# Robust acidic pH and digestive enzyme stability of anti-SarsCov2 IgY antibodies: Implications for treatment of viral transmission thru gastrointestinal, ocular, nasal and skin tissues

**DOI:** 10.1101/2023.08.29.555356

**Authors:** Sanjana Battula, K Saranya, Kranti Meher, R.N Arpitha Reddy, Gopi Kadiyala, Subramanian Iyer, Subrahmanyam Vangala, Satish Chandran, Uday Saxena

**Affiliations:** Reagene Innovations Pvt Ltd; Kyntox Bio Pvt Ltd; Prodigy Inc

## Abstract

There are many paths for the transmission of SarsCov2 virus. The main routes are nasal and oral and the droplets carrying the virus can also be transmitted thru ocular and skin tissues. The gastrointestinal (small intestine), nasal, ocular and skin tissues all present an acidic pH milieu and therefore any treatment with antibodies thru these routes has to have the antibodies remain active at acid pH as well as be resistant to typical protease digestion.

To this end we profiled our anti-SarasCov2 receptor binding domain IgY antibodies for retention of activity at acidic and basic pHs and trypsin digestion. We find that the IgY are strongly resistant to denaturation at acid pH as well as not digested by trypsin. Our data strongly support the use of these IgY in treatment of viral transmission thru the GI, nasal, ocular and skin all tissues where the pH is acidic. We also provide an enabling platform to rapidly asses the suitability of any antibody or protein therapeutic for use at acidic pH.

## Introduction

SarsCov2 virus continues to plague the world with persistent infections thru its many mutant versions. While the vaccines developed have helped, yet the new variants of omicron mutations are still active and often escape the vaccine generated neutralizing antibodies. Thus, there remains a continued need for therapeutic approaches that can block the infections.

We have developed chicken IgY against SarsCov2 receptor binding domain (RBD) which neutralize the binding of virus RBD to ACE2 (the human cell receptor). The IgY were effective in blocking the binding of Wuhan, delta and omicron strain RBD to ACE2 suggesting a broad array of neutralizing capability. Furthermore, the antibodies also reduced the infectivity of human cells by delta strain live virus in vitro. The IgY have been formulated as a drink (the IgY containing egg powder is dissolved in water and the drink is called ImmunIgY) to be used as a swish and swallow passive immunization approach to neutralize the virus in oropharyngeal cavities and GI.

While the nasal and oral routes are considered the most prolific routes of infection, it’s also been known that droplets containing the virus can also enter thru the ocular and skin tissues. Therefore, for complete protection against infection, it is ideal to be able to deliver to antibodies via all the routes.

Within the GI tract, it has been shown that the small intestine is the tissue with the highest expression of ACE2, even higher than lung, and therefore a high sensitivity to infection is present in the gastrointestinal (GI) tracts (1,2). Additionally, ACE2 receptors are present in the nose, mouth, stratified epithelial cells in the upper oesophagus, and absorbent enterocytes of the ileum and colon as well as skin. GI symptoms are present in approximately 50% of COVID-19 patients, consisting of diarrhoea, nausea, vomiting, and abdominal discomfort. Considering the overall evidence, several reports have expressed concerns of possible faecal-oral transmission of SARS-CoV-2 (3,4,). These findings are not unexpected as other well-known coronaviruses, including those causing the MERS and SARS epidemics, have shown to infect enterocytes, leading to several GI symptoms.

It has been reported that the nasal, ocular, skin and of course the GI maintain an acidic environment. In addition, the GI has digestive enzyme trypsin in the small intestine which can also destroy any antibody directed towards the virus. We therefore performed a systematic evaluation of the SarsCov2 IgY that we developed against a range of pH from acidic to basic as well as susceptibility to trypsin digestion. We report here that the IgY or the egg powder containing IgY’s exposure to acid pH as well as trypsin digestion strongly retained the neutralising property of blocking RBD binding to ACE2. These data strongly support the utility of the IgY for blocking infection thru GI, nasal, ocular and skin tissues.

## Methods

### ELISA Protocol

The ELISA protocol reported here is similar to that used by us previously (5). Essentially the steps were as follows:

1. Coat a 96-well ELISA plate with 100 μL of prepared IgY drink.
2. Expose the plate to different pH buffers – 1,2,3,4,5,7 or to trypsin digestion
3. Keep the plate at 4°C overnight.
4. Wash the plate with 400 μL of wash buffer(PBST) three times.
5. Add 200 μL of BSA (PBS + 1% BSA solution) to all the wells.
6. Incubate the plate at room temperature for 1 hour.
7. Wash the plate with 400 μL of wash buffer(PBST) three times.
8. Add 100 μL of B-RBD (x μg/mL) to all the wells.
9. Incubate for 2 hours at 37°C.
10. Wash the plate with 400 μL of wash buffer(PBST) three times.
11. Add 50 μL of HRP conjugate to all the wells.
12. Incubate for 30 mins at 37°C.
13. Wash the plate with 400 μL of wash buffer(PBST) three times.
14. Add 100 μL of TMB solution to all the wells.
15. Incubate for 15 mins at room temperature (dark).
16. Add 50 μL of stop solution to all the wells
17. Read OD at 450 nm.

### Haemagglutination assay (HA)

The HA we used was that reported by others (6), with the major difference being that the assay was performed using a dot blot style assay on filter paper and not in the V-bottom plates. RBCs from chicken blood were used and the degree of agglutination was quantitated by ImageJ analysis.

## Results

### 1. pH effect on binding of anti-SarsCov2 RDB IgY containing egg powder to RDB in ELISA assay

We first tested whether the incubation of IgY containing egg powder with buffers of various pH (pH 1-9) would have an impact on its binding ability to Wuhan RBD using and ELISA method. As shown in Figure 1, most of the pH whether acidic or basic did not impair the IgY’s ability to bind to RBD. It is especially relevant to note that at pH 3 and 4 the binding was the highest suggesting its resistance to acid pH and potential use in pH milieu tissues.

**FIGURE 1.**
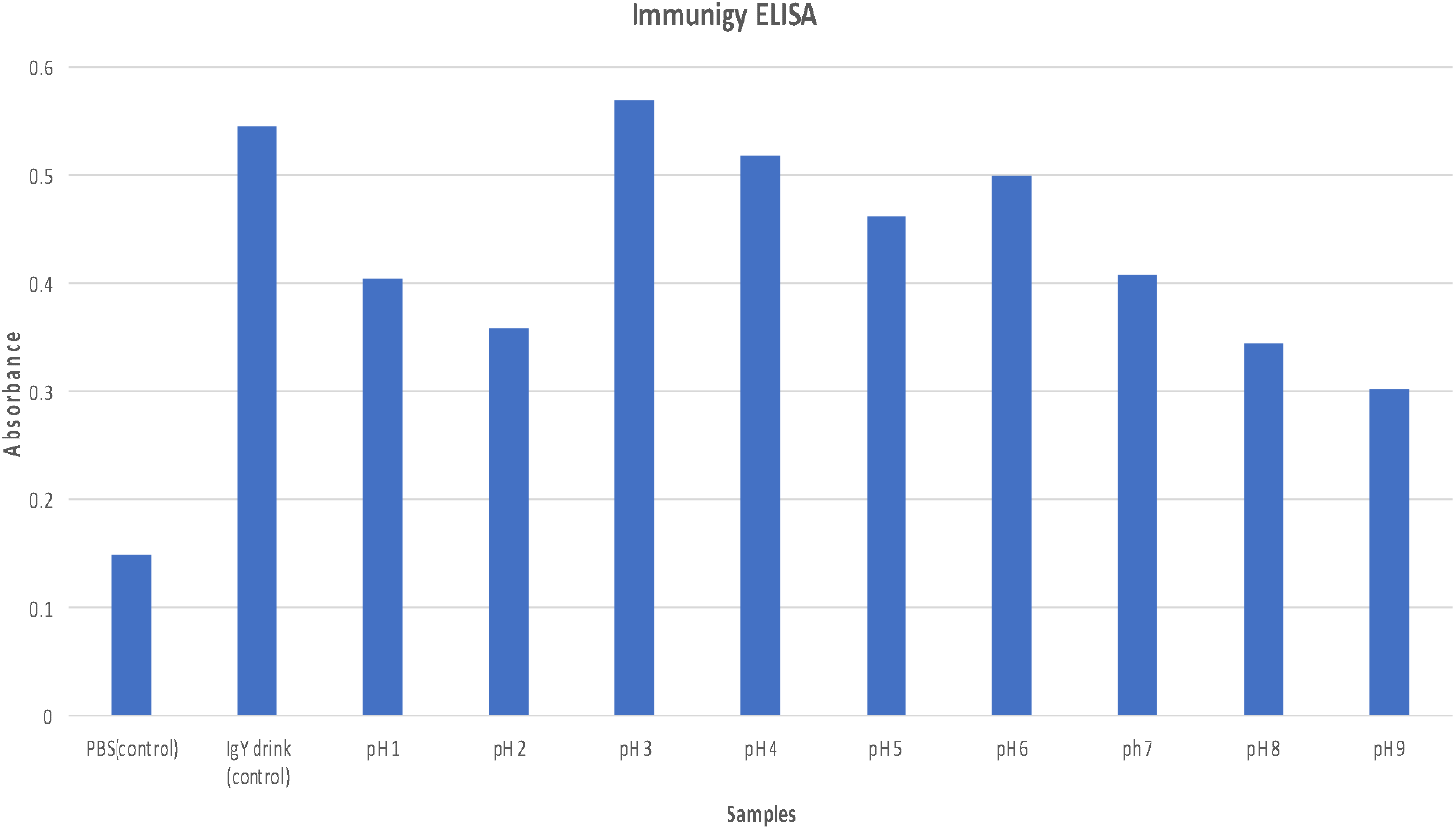

### 2. pH effect on neutralisation activity of anti-SARS-CoV-2 2 IgY antibodies

We then tested if the pH had any effect on the ability of IgY to neutralize the binding of RBD to ACE2 using a dot blot technique. Similar conditions to the one used in ELISA were used and then the IgY tested for neutralising activity. As shown in figure 2 we found that like the ELISA result, exposure to a range of pH from 1-9, the neutralising activity was retained thru the range suggesting that acidic pH incubation do not impar IgY neutralising activity. Further extension of these data suggest that the IgY will be active in tissues with acidic pH

**FIGURE 2.**
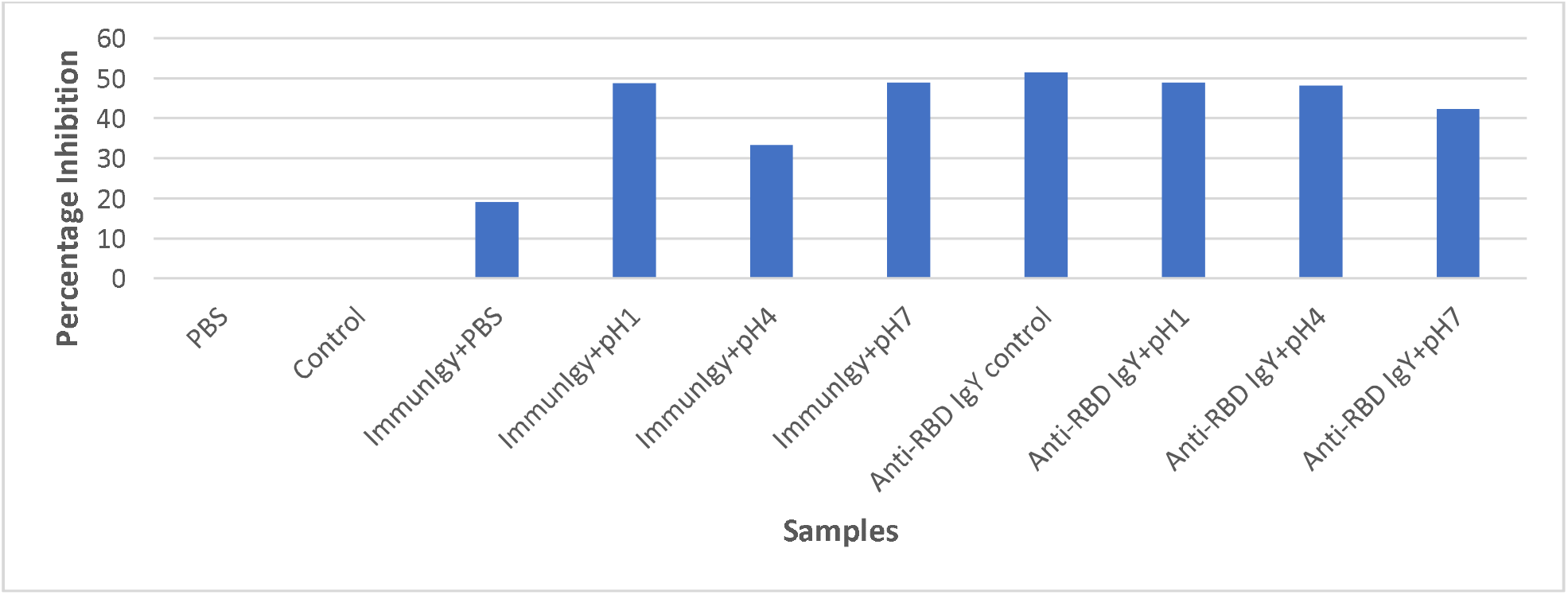

### 3. Food component effect on binding and neutralising activity

Transmission thru the intestine remains a major route for the virus especially in situations where the drinking water may be contaminated as well as the faecal-oral transmission due to poor sewage treatment. To further test any food effect on IgY binding to RBD (via ELISA) and its neutralising activity (dot blot) we performed a series of experiments by testing effect of three major food components separately i.e., protein (in form of BSA), carbohydrate (in form of sucrose) and fat (in the form of vegetable oil) at the pH range used above.

Figures 3-5 below show the impact of each of these food components in ELISA and companion neutralising activity. As shown in Figure 3 there was some diminution of binding by protein in the ELISA assay, not at pH1 but more at higher pH

**FIGURE 3.**
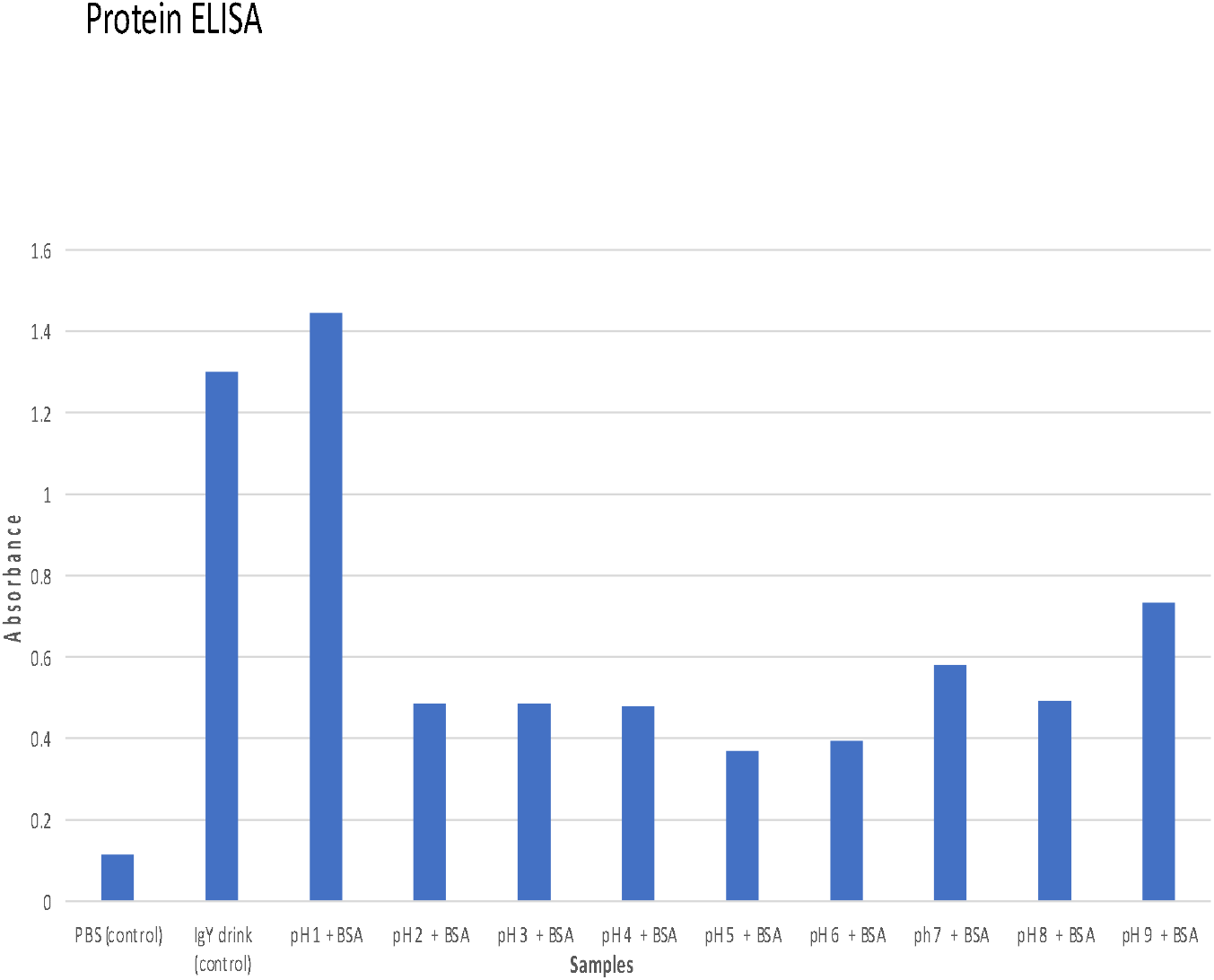

**FIGURE 4.**
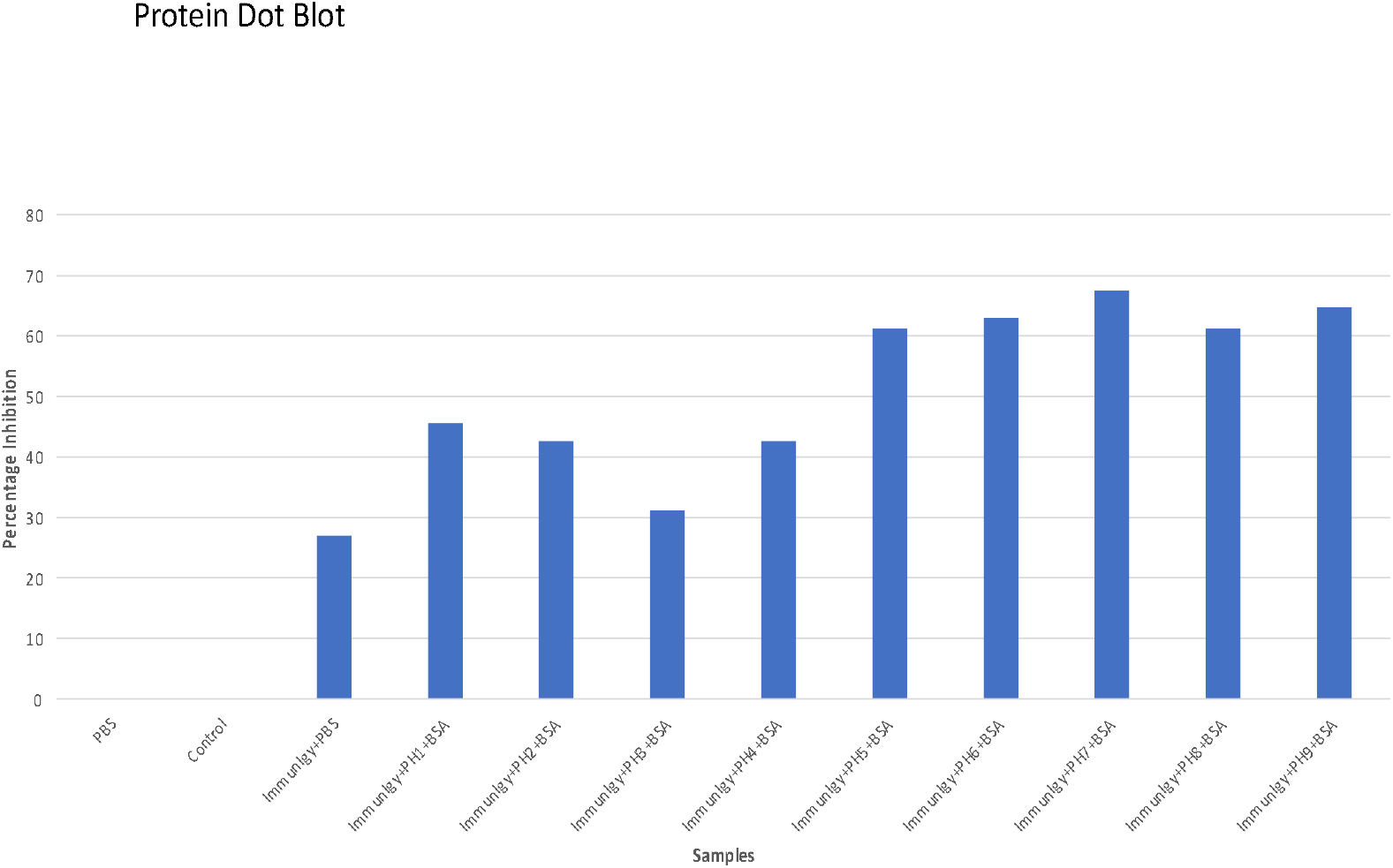

**FIGURES 5.**
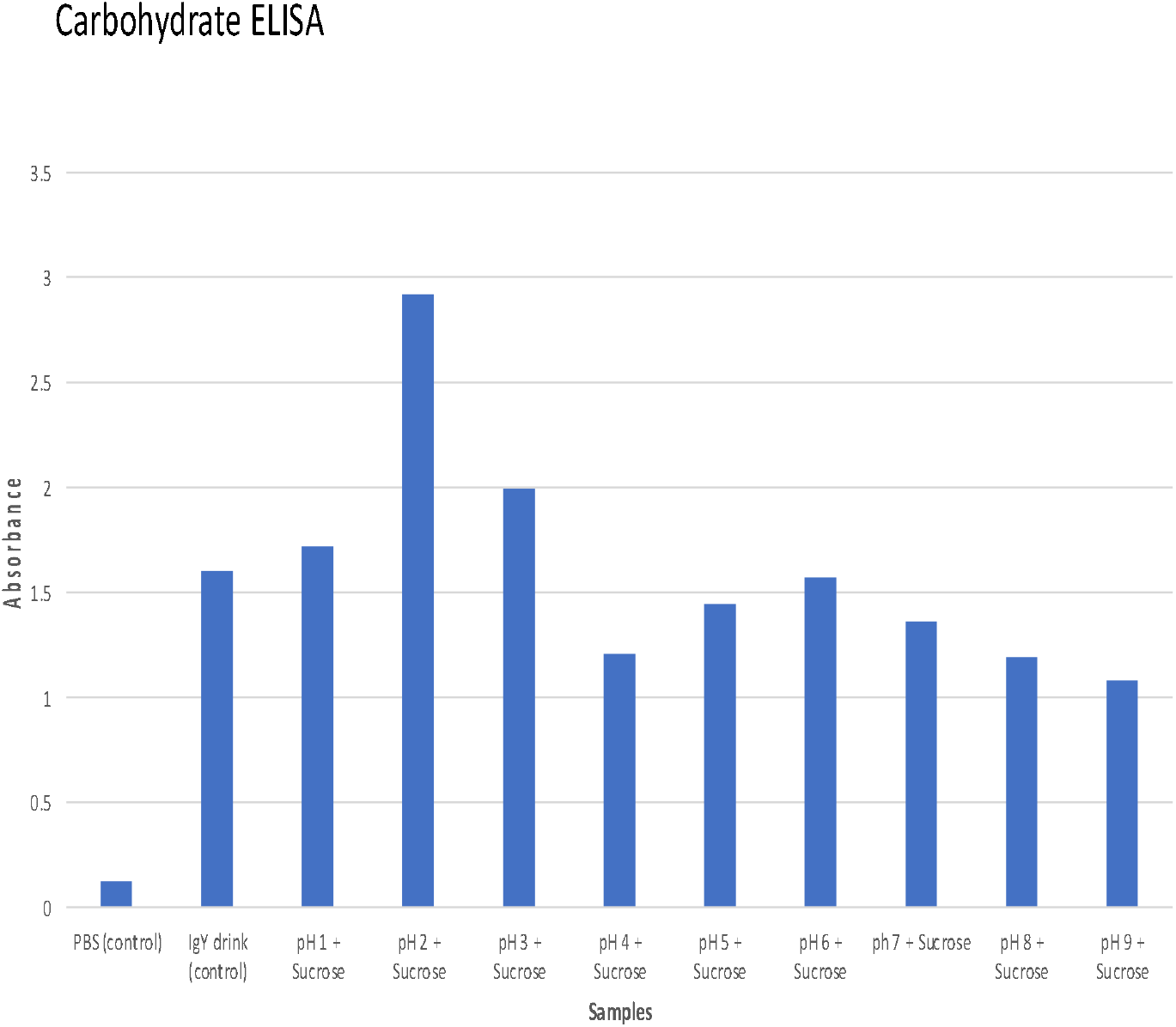

We the performed a companion neutralising activity study. As shown in figure 4 there was no reduction relative to the original ImmunIgY in neutralising activity. These data show that proteins in food are unlikely to have any impact on IgY neutralising activity

We next explored the same with carbohydrate as the food component effect. As shown below in both ELISA and dot blot (figures 5 and 6) there was not much of an effect of carbohydrate especially on neutralising activity of ImmunIgY.

**FIGURES 6.**
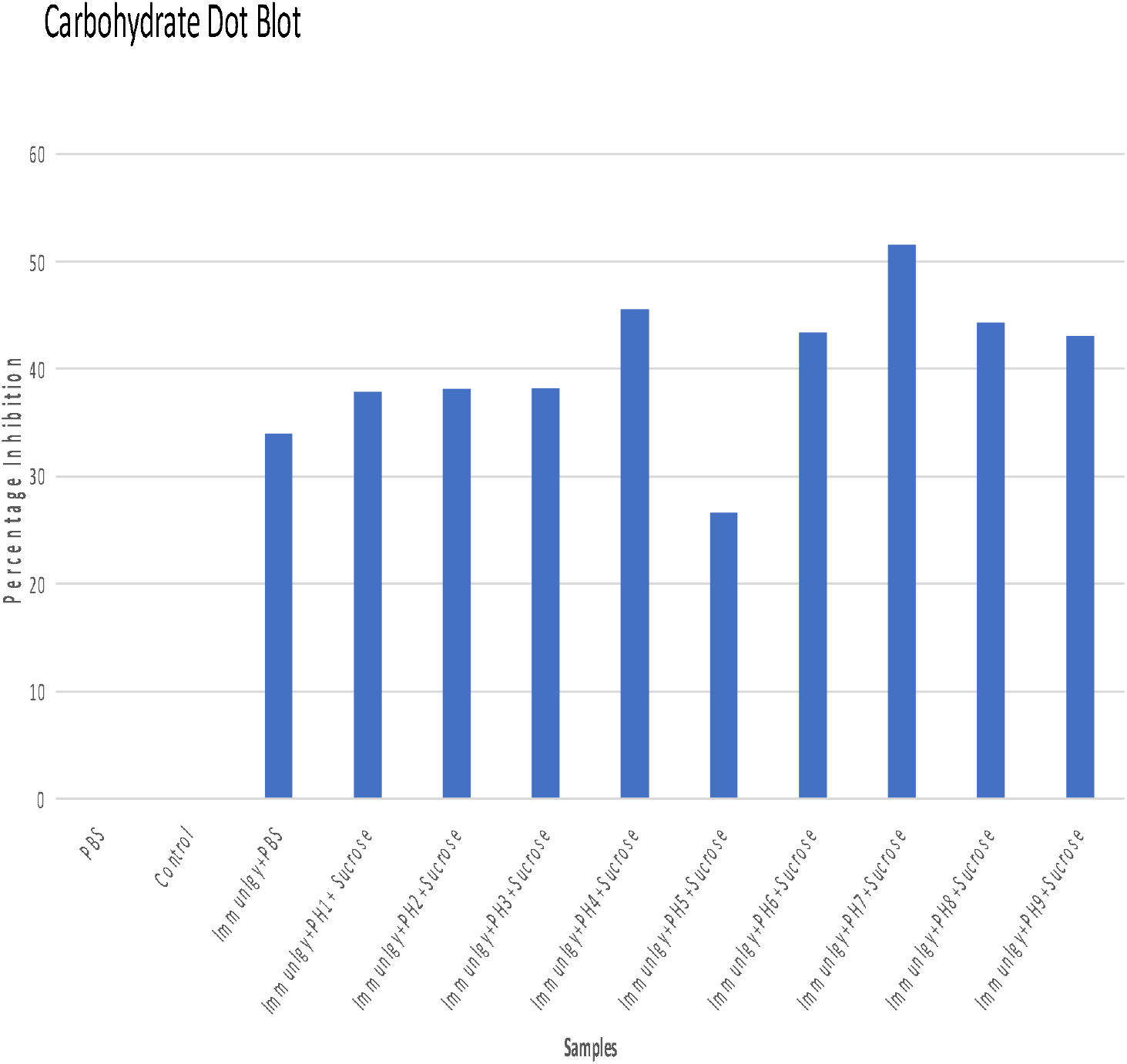

We continued to explore the effect of fat as a food component on both IgY binding to RBD and neutralising activity. Vegetable oil was used as the source of fat. As shown below in Figures 7 and 8. There were only modest changes in the binding or neutralizing activity of IgY in the presence of fat relative to the drink alone.

**FIGURE 7.**
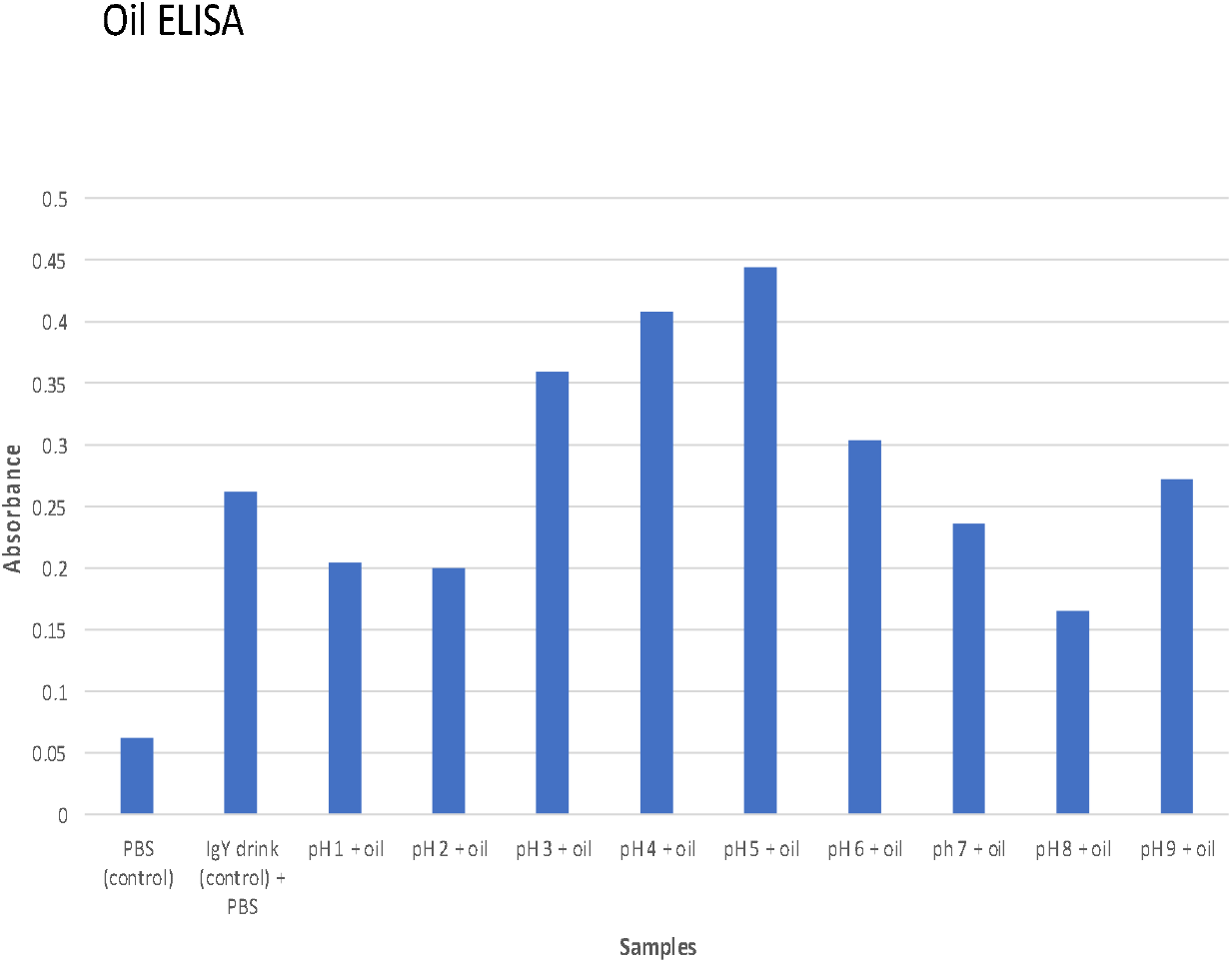

**FIGURE 8.**
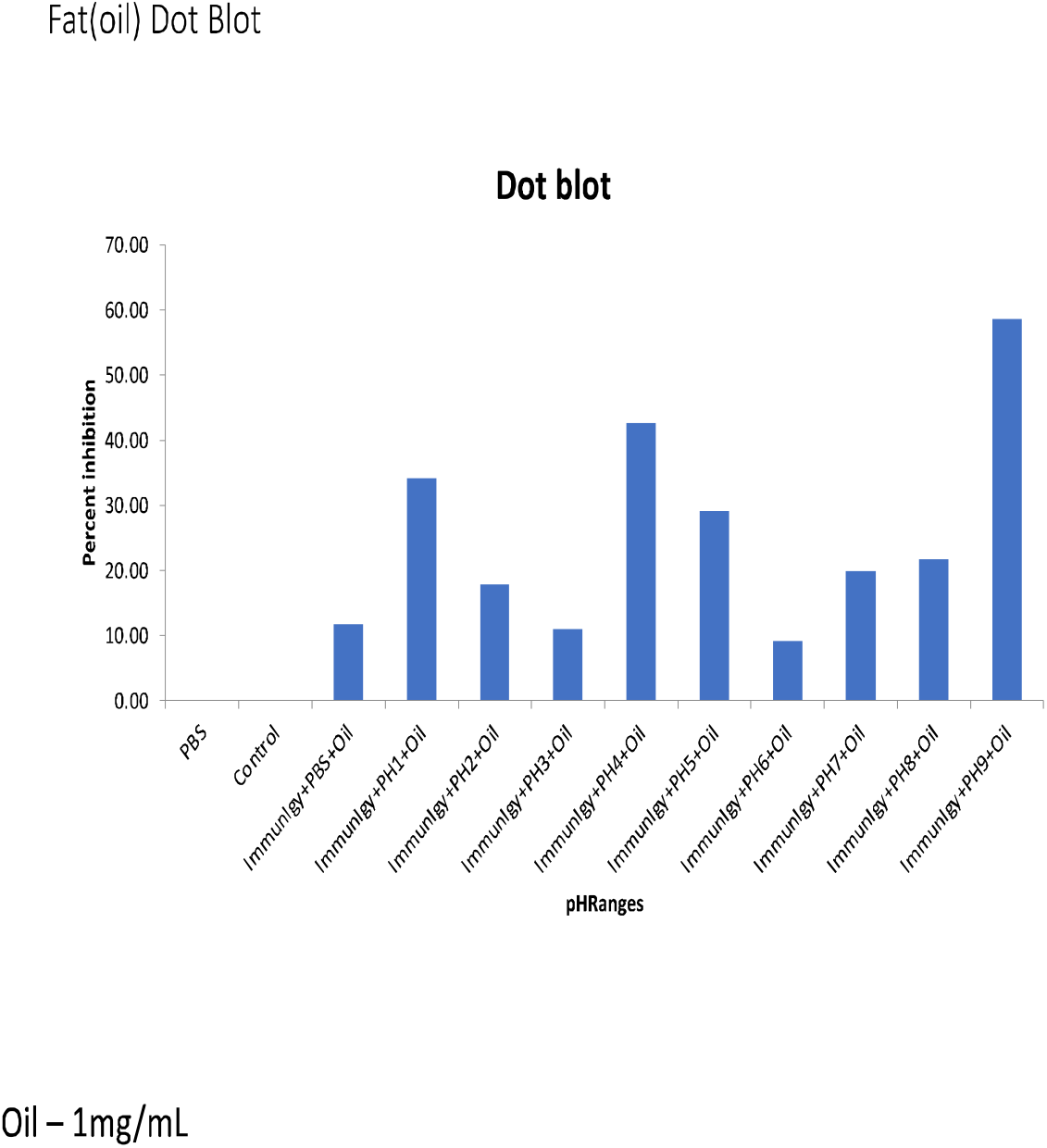

### 4. Effect of trypsin digestion on neutralising activity

There are two major hinderances to the activity of antibodies/proteins in the GI tract, one is the acid pH environment and the other is digestions by trypsin especially in the small intestine which is the major site of GI SarsCov2 viral entry. Our studies presented above show that IgY retains activity at acidic pH so we next tested resistance to enzymatic breakdown by trypsin. Shown below is an SDS-PAGE (figure 9) evaluation of purified anti-SarsCov2 IgY after incubation with trypsin after various time points.

**FIGURE 9.**
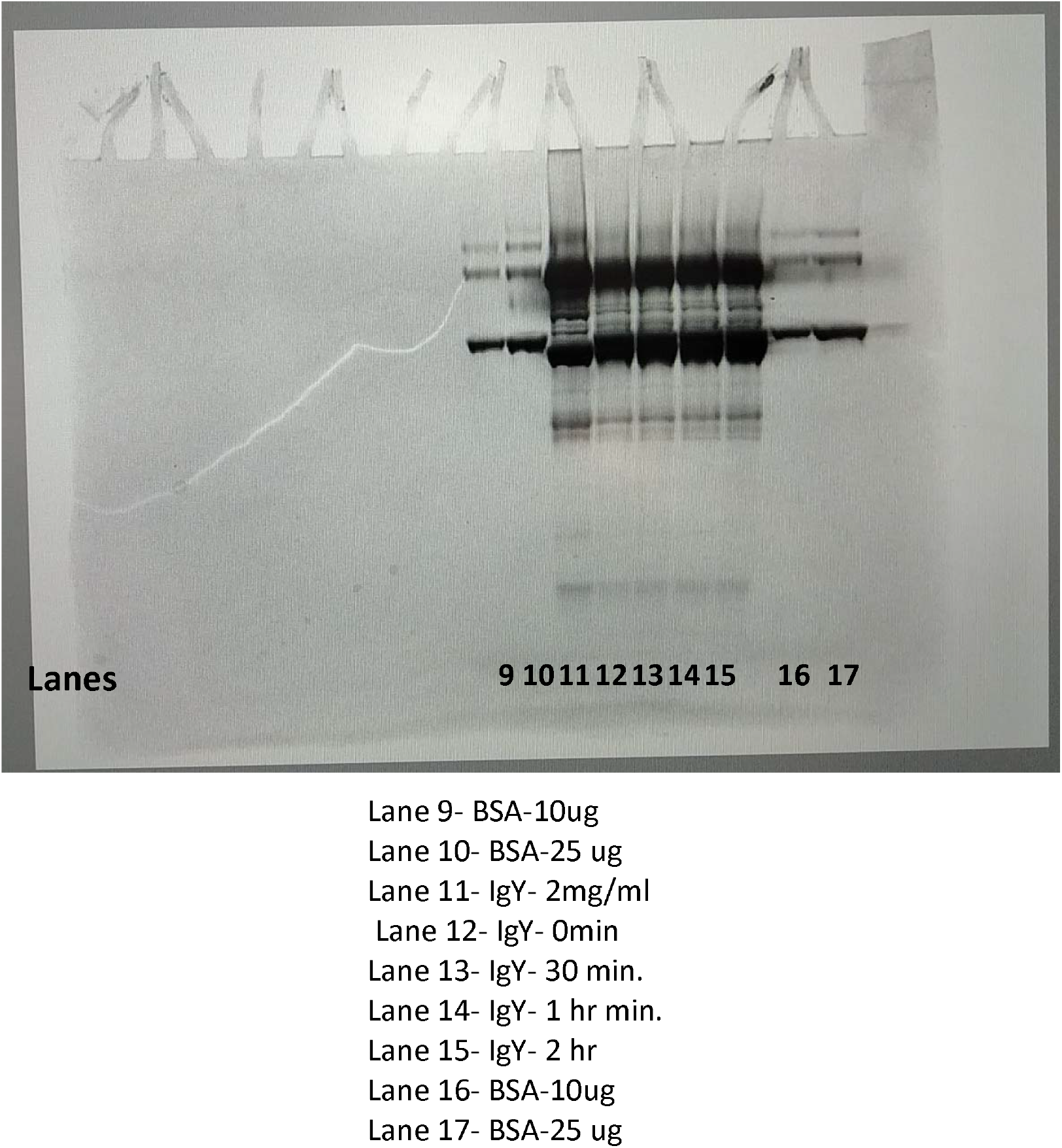

Comparing the protein band intensities across the gel, it is obvious that at any time point of incubation of trypsin, there was no sign of tryptic digestion, since the major bands did not disappear and nor was their appearance of lower band. This suggest that both IgY are resistant to trypsin and would survive in the small intestine.

We then explored if the trypsin digested IgY retained it neutralising activity using dot blot. As shown below (Figure 10) there was no change in activity from base line (time zero incubation with trypsin versus higher incubation times) suggesting that trypsin does not affect the neutralising activity of anti-SarsCov2 RBD IgY.

**FIGURE 10.**
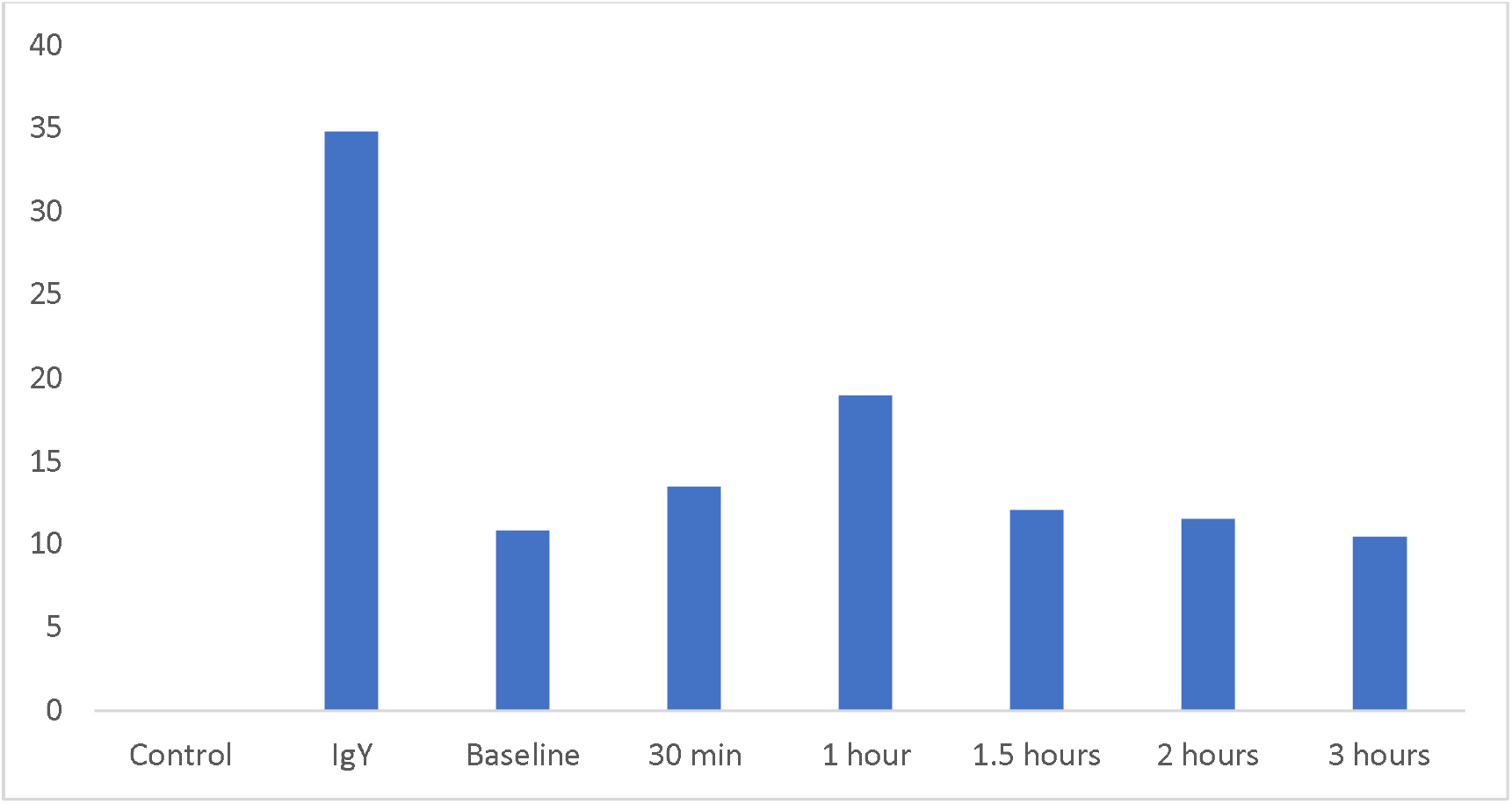

To ascertain if the pH and trypsin resistance are specific to anti-SarsCov2 IgY or is common amongst other IgY we tested pH and trypsin resistance against another IgY raised against an influenza virus antigen. Shown below are the trypsin degradation profile and neutralising activity in a hemagglutination assay which is a standard assay to evaluate anti-influenza antibodies.

**FIGURE 11.**
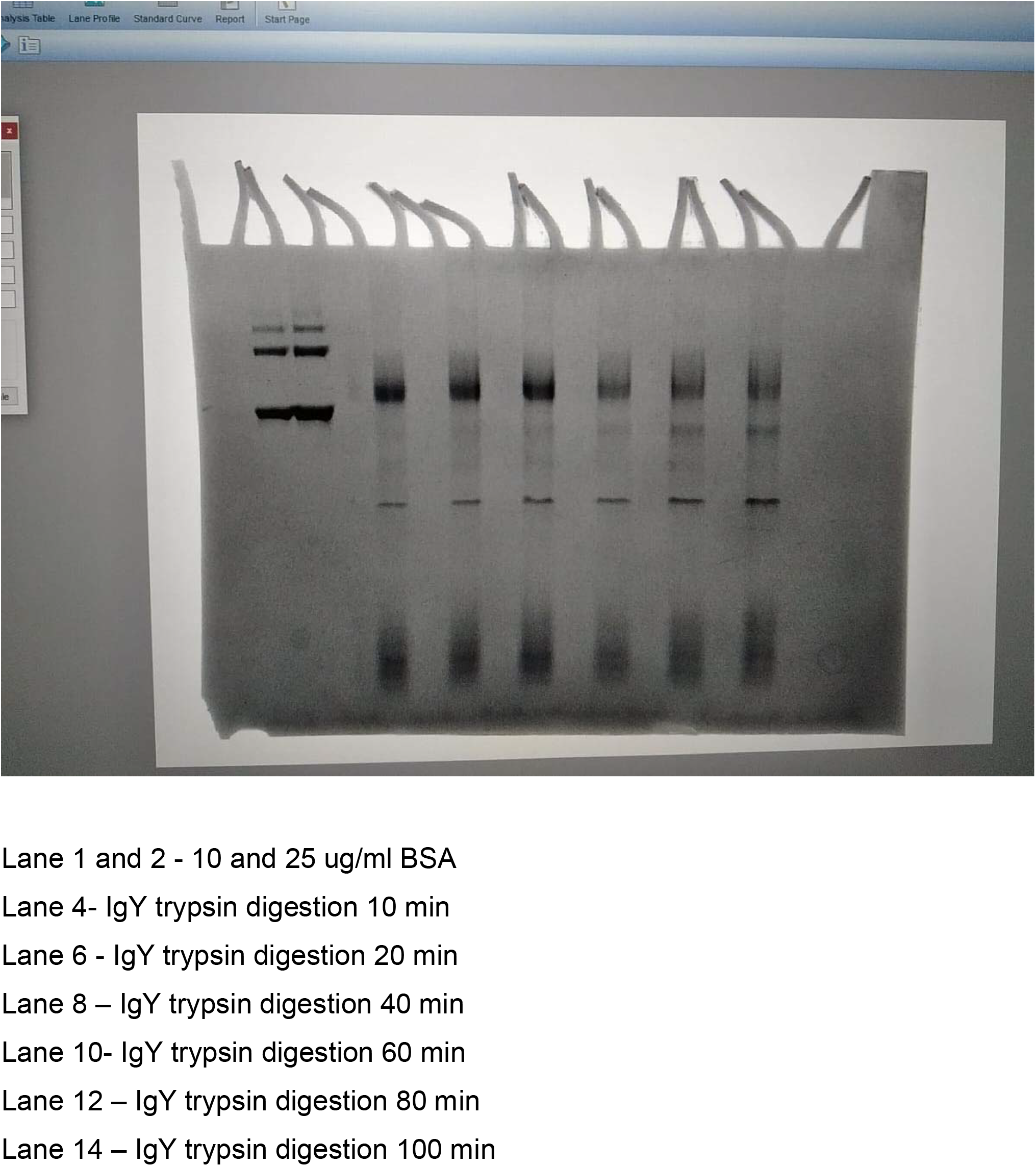

From the SDS-PAGE gel picture above, it is fairly obvious that at time points of 60,80 and 100 min there is degradation apparent from reduction in intensity of higher MW band and appearance of lower MW bands indicative of digestion. This suggest that trypsin resistance is specific to anti SarsCov2 -RBD IgY while another IgY is degraded by trypsin in a time dependent manner.

We then explored if pH affected the functionality of the anti-influenza IgY antibodies using a standard haemagglutination assay. As shown in Figure 10, exposure of anti-influenza IgY to various pH shows that acidic pH from 1-6 markedly decreased the functional activity. These data suggest that the anti-influenza IgY unlike anti-Sarscov2 RBD IgY are susceptible to acid pH and trypsin digestion.

**FIGURE 12.**
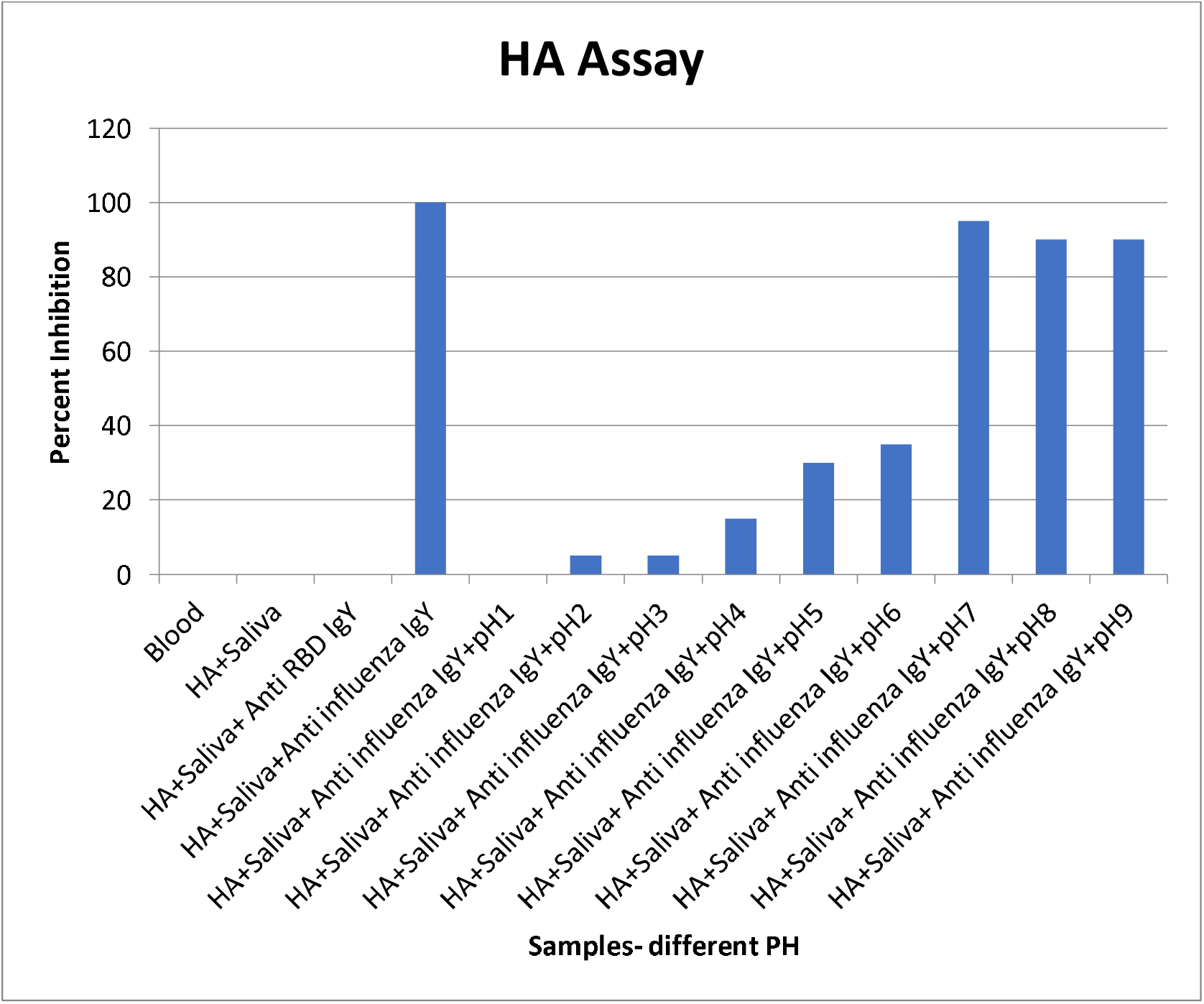

## Discussion

We show here that

1. Anti-SarsCov2 RBD IgY or egg powder containing the IgY are resistant to acid pH denaturation as well as trypsin digestion
2. These data support the development of these IgY as treatment for GI tract infection, use in ocular delivery where pH is acidic, use as nasal drops where too the pH is acidic and finally as skin ointments where the pH is around 4.5
3. These findings are specific to anti-SarsCov2 RBD IgY since another IgY directed towards influenza virus are susceptible to trypsin and acid pH

A classic example of how not sall proteins are suitable for GI delivery is insulin. For decades there have been attempts to deliver it orally without much success. One of the reasons for this failure could be that insulin rapidly degrades at acidic pH. At pH below 5, deamidation at amino acid 21 occurs and covalent dimerization dominates whereas at above pH 8, cysteine disulfide reactions lead to covalent polymerisation of A and B chains of insulin.

Our data also suggest that we can use these studies as an enabling platform to rapidly asses the suitability of antibodies, proteins and peptides for use in GI, skin, ocular and nasal tissues where the pH is predominantly acidic and the GI (small intestine possess trypsin, an enzyme which rapidly degrades mots proteins into amino acids or smaller peptides which can then be absorbed by it. We propose the following steps shown below for this assessment which can be applied to any protein etc. This wet lab process can be preceded by using AI/ML tools to check if there are any “hot spots” in the proteins sequence which may render it suspectable to pH modifications or trypsin digestion.

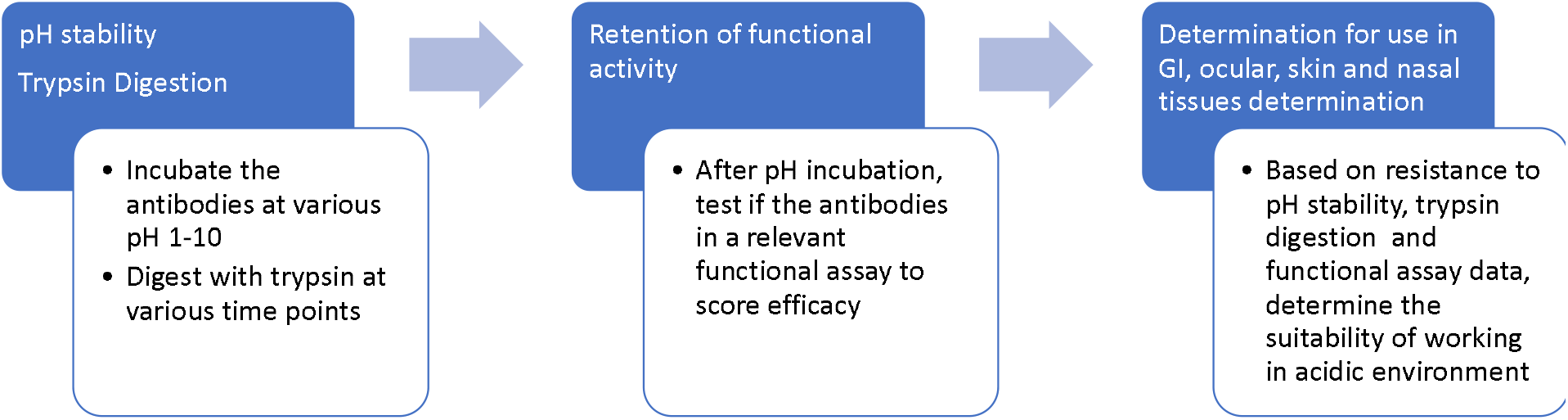
Rapid Method to evaluate acid pH stability and enzyme digestion of antibodies for effectiveness in ocular, GI, skin, nasal tissues

## Acknowledgements

The authors are appreciative of the facilities and support provided at ASPIRE-BioNest, University of Hyderabad, Hyderabad, India.

## Notes

### Competing Interest Statement

The authors have declared no competing interest.

